# Using zebrafish to elucidate glial-vascular interactions during CNS development

**DOI:** 10.1101/2021.01.15.426890

**Authors:** Robyn A. Umans, Carolyn Pollock, William A. Mills, Harald Sontheimer

## Abstract

An emerging area of interest in Neuroscience is the cellular relationship between glia and blood vessels, as many of the presumptive support roles of glia require an association with the vasculature. These interactions are best studied *in vivo* and great strides have been made using mice to longitudinally image glial-vascular interactions. However, these methods are cumbersome for developmental studies, which could benefit from a more accessible system. Zebrafish (*Danio rerio*) are genetically tractable vertebrates, and given their translucency, are readily amenable for daily live imaging studies. We set out to examine whether zebrafish glia have conserved traits with mammalian glia regarding their ability to interact with and maintain the developing brain vasculature. We utilized transgenic zebrafish strains in which *oligodendrocyte transcription factor 2* (*olig2*) and *glial fibrillary acidic protein* (*gfap*) identify different glial populations in the zebrafish brain and document their corresponding relationship with brain blood vessels. Our results demonstrate that *olig2* and *gfap* zebrafish glia have distinct lineages and each interact with brain vessels as previously observed in mouse brain. Additionally, we manipulated these relationships through pharmacological and genetic approaches to distinguish the roles of these cell types during blood vessel development. *olig2* glia use blood vessels as a pathway during their migration and Wnt signaling inhibition decreases their single-cell vessel co-option. By contrast, the ablation of *gfap* glia at the beginning of CNS angiogenesis impairs vessel development through a reduction in Vascular endothelial growth factor (Vegf), supporting a role for *gfap* glia during new brain vessel formation in zebrafish. This data suggests that zebrafish glia, akin to mammalian glia, have different lineages that show diverse interactions with blood vessels, and are a suitable model for elucidating glial-vascular relationships during vertebrate brain development.

## Introduction

Neurons, glia, and vascular cells of the central nervous system (CNS) display unique interactions required for proper brain development and health. Once thought as merely the “glue” that holds the brain together, it has become clear that glial cells are essential to maintain normal brain activity. Similar to how there are different types of highly specialized neurons, glia arise from genetically discrete populations and develop distinct morphology, distributions, and roles. Glia perform a variety of vital functions including supporting synaptogenesis (Eroglu and Barres, 2010), neurotransmitter and ion homeostasis (Murphy-Royal et al., 2017), functional hyperemia (Attwell et al., 2010; MacVicar and Newman, 2015), stabilizing nascent vessels (Ma et al., 2013), and blood-brain barrier (BBB) maintenance (Heithoff et al., 2020).

During development, glia play a critical role in brain maturation, including CNS angiogenesis or the formation of new blood vessels in the CNS, and barriergenesis, the formation of BBB properties (Biswas et al., 2020). A few of the underlying signaling cascades important for glial-vascular interactions have been elucidated yet gaps in knowledge remain (Abbott, 2002). For example, astrocytes have been shown to release Sonic hedgehog (SHH) to promote embryonic BBB formation and integrity (Alvarez et al., 2011). This appears to be recapitulated following stroke, as oxygen-glucose deprived astrocytes promote angiogenesis via SHH secretion (He et al., 2013). Glia can also utilize the vasculature as a pathway during brain development as illustrated by oligodendrocyte precursor cells (OPCs) that require Wnt signaling to migrate along blood vessels in the mammalian brain (Tsai et al., 2016). Whether zebrafish glia also associate with blood vessels or utilize them as a pathway for migration is unknown. Given the increasing interest in utilizing the zebrafish model system in Neuroscience, we set out to examine glial-vascular interactions during zebrafish brain development.

In this study, we utilized transgenic lines that differentially label well-studied glial populations, *oligodendrocyte transcription factor 2* (*olig2)* and *glial fibrillary acidic protein* (*gfap)*. We show that *olig2* OPCs cells utilize vascular networks as a pathway over time and this relationship is mediated through Wnt signaling. Conversely, *gfap* glia associate along vascular walls and are required for normal vessel development as ablation of *gfap* cells causes irregular vessel development and a reduction in Vascular endothelial growth factor (Vegf) expression. Lastly, intracranially implanted human glia reach out to zebrafish blood vessels, supporting a conservation in endogenous signals that maintain glial-vascular interactions in the zebrafish brain. Taken together, these studies substantiate how the zebrafish model can dissect glial-vascular biology during vertebrate brain development and hold promise for uncovering novel mechanisms in glial Neuroscience.

## Methods

### Ethics Statement

All zebrafish husbandry and experiments were done in accordance with Virginia Tech Institutional Animal Care and Use Committee (IACUC) guidelines. The Virginia Tech IACUC abides by the Guide for the Care and Use of Laboratory Animals and is accredited by the Association for Assessment and Accreditation of Laboratory Animal Care (AAALAC International).

### Zebrafish Husbandry and Lines

Adult zebrafish (*Danio rerio*) were maintained according to Virginia Tech IACUC guidelines on a 14/10 hour light/dark cycle in a Tecniplast system at 28.5°C, pH 7.0, and 1000 conductivity. The *Tg(glut1b:mCherry)* line was obtained from Dr. Michael Taylor’s laboratory at the University of Wisconsin-Madison. The *Tg(fli1a:eGFP)*^*y1*^ and *casper* lines were obtained from the Sinnhuber Aquatic Research Laboratory at Oregon State University. The *Tg(olig2:dsRed), Tg(gfap:GFP)*, and *Tg(gfap:nfsB-mCherry)* lines were obtained from Dr. Sarah Kucenas at the University of Virginia. The *Tg(kdrl:mCherry)* line was obtained from Dr. Wilson Clements from St. Jude Children’s Research Hospital. Fixed *Tg(flk1:mCherry)* embryos were obtained from Dr. Chris Lassiter at Roanoke College. Animals were sorted for fluorescence and brain microbleeds on an upright Nikon SMZ18 stereomicroscope, courtesy of the Pan Laboratory.

For all experiments, embryos were collected from multiple pair-wise crosses. Embryos were maintained in embryo water (1.5g Instant ocean salt per 5L RO water) at 28.5°C. For studies with lines not bred onto the *casper* background, 24 hour post-fertilization embryos were treated with 200µM N-Phenylthiourea (PTU) (Sigma # P7629) to prevent melanocyte formation. Larvae for human astrocyte microinjections were acclimated to 32-35°C to accommodate a more desirable temperature for astrocyte cell growth.

### Database Mining

A list of human glial genes was prepared from the database generated by Cahoy et al. (Cahoy et al., 2008). Ensembl was used to determine if a human glial gene was annotated in the zebrafish genome. The description and common name of each gene was validated with GeneCards.

### Reverse Transcription-Polymerase Chain Reaction (RT-PCR)

For RNA extraction and cDNA synthesis, 10 *Tg((fli1a:eGFP)*^*y1*^*;casper)* larvae were collected in RNase-free tubes (Fisher #12-141-368) and humanely euthanized on ice. The egg water was removed and replaced with 50µL of Trizol (Invitrogen #15596026), 5µL of Trizol per larvae. Total genomic RNA was extracted using the PureLink® RNA Mini Kit (Ambion #12183025) adapted with TRIzol™/Chloroform (Invitrogen # 15596026, Fisher #ICN19400290) according to the manufacturer’s guidelines. Samples were eluted in 30µL nuclease-free water. RNA concentration and quality was assessed with a Nanodrop. 1µg of cDNA was synthesized using SuperScript™ VILO™ Master Mix (Invitrogen #11755250) according to manufacturer’s guidelines. 10ng of cDNA was loaded per PCR reaction. *elfa* was used as an internal housekeeping gene as its stability has been noted over developmental time points in zebrafish compared to other controls used in other model systems (McCurley and Callard, 2008). Non-template control reactions were performed for all runs to rule out reagent contamination. Primer sequences, genome annotations, annealing temperatures, and expected product sizes are listed in **Supplemental Table 2**. Samples were loaded on a 1% agarose gel and electrophoresis was performed at 120V for 40 minutes before imaging gels on an Azure Biosystems c500 imager.

### Reverse Transcription-Quantitative Polymerase Chain Reaction (RT-qPCR)

For cDNA template generation, 20 *Tg(kdrl:mCherry)* larvae were collected per tube in 50µL of TRIzol™, in quadruplicate, per time point from 0 to 7dpf. RNA extraction and cDNA synthesis was performed as described above. For RT-qPCR reaction set-up, 50ng of cDNA template was loaded per well, ran in technical triplicates, with 3-4 biological samples per time point. TaqMan™ Universal Master Mix II, no UNG (Thermo # 4440040) was utilized according to manufacturer’s guidelines for amplification with Taqman assays against glial genes and an internal control (*elfa/eef1a1l1*). The following Assay IDs were used: *aqp4* (Assay ID: Dr03095840_m1), *glt-1/slc1a2b* (Assay ID: Dr03119707_m1), *glula* (Assay ID: Dr03433569_g1), *gfap* (Assay ID: Dr03079978_m1), *glut1b/slc2a1a* (Assay ID: Dr03103605_m1), and *eef1a1l1* (Assay ID: Dr03432748_m1). Custom primers were generated for a few targets with the following Assay ID and Zebrafish Ensembl Transcript ID: *kir4*.*1*/*kjcn10a* (Custom Assay ID: ARXGT6V, Ensembl: ENSDART00000172874.2), *aldh1l1* (Custom Assay ID: ARU64ZZ, Ensembl: ENSDART00000151205.3), and *glast/slc1a3b* (Custom Assay ID: ARMFY6E, Ensembl: ENSDART00000131693.3). Glial genes were amplified with the FAM-MGB reporter dye and *eef1a1l1* was amplified with a VIC-MGB_PL reporter dye. An Applied Biosystems StepOnePlus™ Real-Time PCR System was used for RT-qPCR amplification and the standard program according to manufacturer’s guidelines ran for 40 cycles. Delta C_T_ (ΔC_T_ or 2^-CT^) was calculated according to standard protocol, whereby the glial gene data is presented relative to *eef1a1l1* control expression (Schmittgen and Livak, 2008).

### Pharmacological Treatments

10mM Metronidazole (MTZ) (Sigma #M3761-5G) was used for *Tg(gfap:nfsB-mCherry)* ablation studies as previously reported (Johnson et al., 2016). For MTZ studies, animals were not dechorionated at 8 hours post-fertilization (hpf) to prevent damage to the developing tissue. MTZ was dissolved in a 0.2% Dimethyl sulfoxide (DMSO) (Sigma #D8418)/PTU solution. To prevent any light-mediated degradation, MTZ was made in amber tubes and the multi-well plates containing treated animals were wrapped in aluminum foil. For Wnt and Vegf small molecule studies, XAV939 (Cayman Chemical #13596) and Tivozanib (AV951) (Apexbio Tech #A2251) were re-suspended as10mM stocks in DMSO and used at final working concentrations of 30µM XAV939 and 1µM AV951.

### Live Zebrafish and Fixed Tissue Confocal Microscopy

Live animals were removed from multi-well plates with a glass Pasteur pipette and anesthetized with 0.04% MS-222 (Sigma #A5040) in individual 35mm petri dishes containing a 14mm glass coverslip (MatTek Corporation #P35G-1.5-14-C). A 1.2% low melting point agarose (ThermoFisher #16520050) solution was made in embryo water for embedding. MS-222 was removed from the animal and a bolus of warm agarose the size of the coverslip was added to the animal/cover glass. A dissection probe was used to align animals dorsal side down for brain imaging. After the agarose solidified, the petri dish was filled with 0.04% MS-222 (live animals) or phosphate buffered saline (PBS) (Fisher # BP661-10) (fixed animals) and the dish was wrapped with parafilm along the edges. Confocal microscopy was performed to take z-stacks at optimal section thickness with an Olympus Fluoview FV1000 confocal microscope and a 40X(NA0.75) objective. The Nyquist optimal section thickness value for z-stacks was calculated through an algorithm in the Olympus Fluoview software for all images. Because this is an upright microscope, the glass coverslip bottom dishes containing the animals were flipped upside down so that the glass side made contact with the objective.

For fixed whole mount tissue, Vegfa staining was captured on the Olympus F1000 with a 40X (NA 0.75) objective. For Gfap staining, embryos were imaged on a Nikon A1R confocal microscope with a 10X(NA 0.45) objective.

### Live Multiphoton Microscopy (MPE)

Live multiphoton microscopy was performed to take z-stacks at a 1µm optical section thickness with a four-channel Olympus FV1000MPE multiphoton laser scanning fluorescence microscope equipped with a mode-locked Coherent Chameleon two-photon laser and XLPLN25X/1.05NA Olympus water-immersion objective. Volumetric reconstructions and orthogonal views were created using NIS Elements software. Line plots were generated using the Olympus Fluoview software.

### Astrocyte culture and Immunocytochemistry (ICC)

Human astrocyte cultures (ScienCell #1800) were maintained at 10% CO_2_ in specialty media (ScienCell #1801). For Green Fluorescent Protein (GFP) expression, cells were transduced (MOI=2) with a lentivirus constitutively expressing GFP (OriGene Tech # PS100093V). A pure GFP population was generated with puromycin selection for a few days in culture before use in experiments and cell line storage.

For ICC of differentiated astrocyte markers, roughly 200,000 cells were seeded onto 12mm glass coverslips in a 6-well plate. After adhering overnight, cells were rinsed in PBS and then fixed for 10 minutes at room temperature in 4% PFA. Coverslips were rinsed in PBS, washed in PBST (PBS+0.1% Tween20) several times, and then blocked in 10% Normal Donkey Serum made in PBST for one hour at room temperature. After blocking, coverslips incubated in the following primary antibodies; chicken anti-GFAP (Abcam #ab4674,1:1,000), rabbit anti-Kir4.1(Alomone #APC-035, 1:1,000), mouse anti-ALDH1L1(UC Davis NeuroMab #75-164 1:500), and guinea pig anti-GLT-1(Millipore #AB1783 1:1,000) overnight at 4°C. Coverslips were washed in PBST for 30 minutes, and incubated with secondary antibodies; goat anti-chicken-Alexa 647, goat anti-rabbit-Cy3-550, goat anti-mouse-Cy3-550, and goat anti-guinea pig-Alexa 647 (Jackson Immuno Research, 1:1,000) at room temperature for 1 hour. Coverslips were counterstained with DAPI (ThermoFisher #D1306, 1:1000) for 5 minutes and washed in PBST for 25 minutes before mounting onto charged slides (VWR #48311-703) with Fluoromount (Electron Microscopy Sciences #17984-25). Images were acquired with a 40X objective (NA 0.75) and a 2.0X optical zoom on an Olympus FV1000 confocal microscope.

### Astrocyte microinjection

Cultured human astrocytes labeled with GFP were counted according to manufacturer’s guidelines and re-suspended at 25,000cells/µL in PBS. 6 dpf *Tg(kdrl:mCherry)* larvae were anesthetized with 0.04% MS-222 (Sigma #A5040) prior to microinjection. During anesthesia, thin wall glass capillary needles with filament (World Precision Instruments #TW150F-4) were pulled with program 4 “Pro-Nuclear Injection” (Heat: 460, Pull:90, Vel: 70, Delay: 70, Pressure: 200, Ramp: 485) on a horizontal pipette puller (Sutter, Model P-1000). This program creates needles with a 0.7µM tip and a taper of 6-7mm. 1µL of Phenol Red (Sigma #P0290) was added to 9µL of the 25,000 cells/µL solution mixture. Capillaries were positioned in a round glass pipette holder and back loaded with 1uL of cell mixture. Each loaded needle was calibrated using a pneumatic pump and micrometer (Carolina Biological Supply #591430) to calculate the bolus and corresponding cell number per injection. Animals were positioned dorsal side up for intracranial injection in a homemade agarose injection mold, and then microinjected with around 25 cells. While we attempted to only implant 25 cells (1nL) per animal, microinjections are not trivial and slight pressure differences between animals could lead to more or less cells implanted. After injections, animals recovered and were tracked in individual wells of a 24-well plate 24 and 48 hours later.

### Olig2-Fli1 Interaction and single cell analyses

Imaging started at the anterior mesencephalic central artery and ended at the posterior mesencephalic central artery. Interactions between o*lig2* glia (red) and *fli1* vessels (green) were quantified within 130µm of the zebrafish brain using the orthogonal view in Olympus Fluoview software. For single *olig2* glia cell counts, a threshold was created for the bottom and top half of the image due to the top half being brighter. Thresholds were set according to the control group for each data set. OIB images were saved as TIFFs and quantified with the 3D bright spot feature in NIS Elements software.

### Glut1 fluorescence analyses

Confocal stacks were opened in FIJI-ImageJ analysis software and the *mCherry* fluorescent signal was measured through the Integrated Density feature for each image.

### Vessel distance analyses

Vessel height and diameter of the distance between the basal communicating artery (BCA) and posterior communicating segments (PCS) was measured using ImageJ. The OIB file for each image was opened in ImageJ, and measurements for height were taken on the left side of the brain by drawing a line segment from the BCA to the PCS. The length of this line segment was measured using ImageJ. To measure the diameter of the PCS, a line segment was drawn between both ends of the PCS and measured using ImageJ.

### Western blot

8-10 larvae were pooled per sample for each group, humanely euthanized in a 1.5mL tube on ice, and egg water was replaced with 50µL of RIPA lysis buffer. Samples were homogenized with a Pellet Pestle Mixer for 30 seconds on ice. Lysates were then centrifuged at 12,000 rpm for 15 minutes at 4°C. The supernatant was collected and diluted 1:6 for BCA protein quantification (ThermoFisher #23227) and performed according to manufacturer’s guidelines. 15µg of each sample was loaded per lane of a MIDI 4-15% Criterion Precast Midi SDS page gel (BioRad #5671085) and electrophoresis was performed at 120mV for 75 minutes. Samples with transferred to PVDF membranes using the Trans-Blot Turbo Transfer system (BioRad #1704150). Membranes were blocked in blocking buffer (LICOR # 927-80001) for 1 hour at room temperature. Blocked membranes were incubated with a mouse anti-zebrafish VEGF antibody (R&D Systems #MAB1247, 1:1,000) and chicken anti-GAPDH antibody (Sigma #AB2302, 1:5,000) overnight at 4°C. The membrane was washed the next day with TBST (0.1% Tween20) for at least 30 minutes and incubated in blocking buffer with 0.1% Tween 20, 0.01% SDS, IRDye® 800CW Donkey anti-Mouse (Li-COR #925-32212, 1:20,000) and IRDye® 680RD Donkey anti-Chicken (Li-COR #926-68075, 1:20,000) secondary antibodies for 1 hour at room temperature. Membranes were washed again in TBST for at least 30 minutes at room temperature, briefly rinsed in TBS, and were imaged on a Li-COR Odyssey Fc Imager. Densitometry of individual bands was performed in FIJI/Image-J software.

### Zebrafish whole-mount immunohistochemistry (WMIHC)

*Tg(flk1:mCherry)* or *Tg(gfap:nfsB-mCherry)* animals were humanely euthanized at 3dpf in 1.5mL tubes on ice and egg water was replaced with 4% PFA for incubation at 4°C overnight. Larvae were rinsed in PBS to remove PFA the next day. *Tg(flk1:mCherry)* larvae were slowly dehydrated in a PBS/Methanol series and stored in 100% methanol (VWR # BDH1135-1LP) at −20°C until staining. To generate sections for regular immunohistochemistry, larvae were sunk in a 30% sucrose solution overnight at 4°C after removing PFA. Animals were then embedded in an embedding block (EMS # 62534-15) containing O.C.T compound (EMS #62550-0) and flash frozen below a cold alcohol slurry. Blocks were stored at − 80°C until thin 10-12µm serial sections were generated with a Leica cryostat.

For Gfap WMIHC staining, larvae were rehydrated into PBS using a PBS/Methanol series and then rinsed in PBST for 3 × 5 minutes. To permeabilize the tissue, larvae were incubated in Proteinase K (Invitrogen #AM2546,1:800) for 10 minutes at room temperature. PK activity was quenched with 10% normal donkey serum (EMD Millipore #S30-100ML) made in PBST. Larvae were washed 3 × 5 minutes in PBST and then blocked in 10% normal donkey serum for 2 hours at room temperature. A rabbit anti-GFAP (Dako # Z033401-2, 1:250) primary antibody was diluted in blocking solution for incubation at 4°C overnight. Negative control secondary antibody only samples were set aside in blocking solution at the same time. Larvae were rinsed 5 × 5 minutes and then 3 × 20 minutes in PBST. Secondary antibody donkey anti-rabbit Alexa Fluor 488 (Invitrogen #A21206,1:200) was prepared in blocking solution for 4°C overnight incubation. Larvae were then washed for 6 × 15 minutes before proceeding to confocal imaging. For IHC, sections were stained in a humidity chamber with a similar method, excluding PK treatment. Sections were counterstained with DAPI to label nuclei, coverslipped, and then imaged on a Nikon A1R confocal microscope.

To perform Caspase3 staining, larvae were rinsed in PBST (0.03% Tx-100 (Sigma # T9284-500ML)) for 3 × 5 minutes. Antigen retrieval was performed by incubating animals in 150mM HCl (pH 9.0) for 5 minutes at room temperature. The larvae were transferred to a 70°C heat block and incubated for another 10 minutes. Larvae were then washed in PBST for 3 × 5 minutes. To permeabilize the tissue, larvae were incubated in 0.05% Trypsin-EDTA (Thermo Fisher #25300054) on ice for 45 minutes. Two quick PBST rinses were performed for a total of 10 minutes. Larvae were then incubated in a blocking solution of 2% Normal Goat serum (Sigma #S26-LITER), 1% BSA (Fisher#BP1605-100), and 1% DMSO in PBST for 1 hour at room temperature. A rabbit anti-Caspase 3 (BD Pharmingen #559565; 1:500) primary antibody was diluted in blocking solution for incubation at 4°C overnight. Negative control secondary antibody only samples were set aside in blocking solution at the same time. Larvae were rinsed multiple times in PBST for an hour before incubation in secondary antibody, goat anti-rabbit Atto 647N (Sigma #40839-1ML-F, 1:500) overnight at 4°C. Larvae were then washed for 6 × 15 minutes before imaging on an Olympus FV1000 confocal microscope.

### Statistical analyses

For analysis of *olig2*-vessel interactions over time, a one-way ANOVA with Tukey’s multiple comparisons test was performed using GraphPad Prism software to determine p-values. For XAV939 and AV951 small molecule treatment experiments, a one-way ANOVA with Tukey’s multiple comparisons test was performed using GraphPad Prism software to determine p-values. For *glut1* fluorescence analyses, a two-tailed, unpaired student’s t-test was performed using GraphPad Prism software to determine p-values. For Western blot analysis, a one-way ANOVA with Tukey’s multiple comparisons test was performed using GraphPad Prism software to determine p-values. For Caspase3 activation in MTZ-treatment WMIHC studies, a one-way ANOVA with Tukey’s multiple comparisons test was performed using GraphPad Prism software to determine p-values.

## Results

### Olig2 and Gfap characterize distinct glial types in the zebrafish brain

The mammalian CNS contains a number of genetically distinct cell linages. For example, the Olig2 transcription factor characterizes NG2 glial progenitor cells that are destined to become oligodendrocytes (Rowitch, 2004). Astrocytes are characterized by a number of markers, most notably GFAP, Glutamate transporter-1 (*GLT-1*), Aldehyde dehydrogenase 1 family, member L1 (*ALDH1L1*), and Aquaporin 4 (*AQP4*) (Cahoy et al., 2008; Freeman, 2010). While a subset of these lineages interact with the CNS vasculature, the expression of astrocytic markers in parallel with CNS vessel development has not been well studied. We first assessed the presence of commonly expressed mammalian glial genes over critical time points during zebrafish brain development. As shown through RT-PCR analysis in **Figure 1A**, glial genes are developmentally regulated. Most genes were present by 1 day post-fertilization (dpf) and all genes amplified were more robustly expressed by 4dpf. We also amplified the BBB transporter *glucose transporter 1b* (*glut1b/slc2a1a*) to indicate the onset of CNS angiogenesis, as it was previously shown that barriergenesis and CNS angiogenesis occur simultaneously via *glut1b* expression during zebrafish development (Umans et al., 2017). Through follow-up RT-qPCR, we generated a descriptive quantification of fundamental glial markers, (*slc2a1a, slc1a2b, slc1a3b, aldh1l1, aqp4, kcnj10a, gfap*, and *glula*), from 0-7dpf (**Supplemental Figure 1**). RT-qPCR validated the increase of these glial transcripts from 2-7dpf, which correlates with the increase of the CNS vascular-specific *glut1b/slc2a1a* after the maternal to zygotic transition in development. While our PCR panel evaluated only a handful of genes, we also performed database mining in the zebrafish genome for annotated genes that are also highly expressed in human astrocytes (**Supplemental Table 1**) as previously reported in a transcriptomic study (Cahoy et al., 2008).

**Figure 1.**
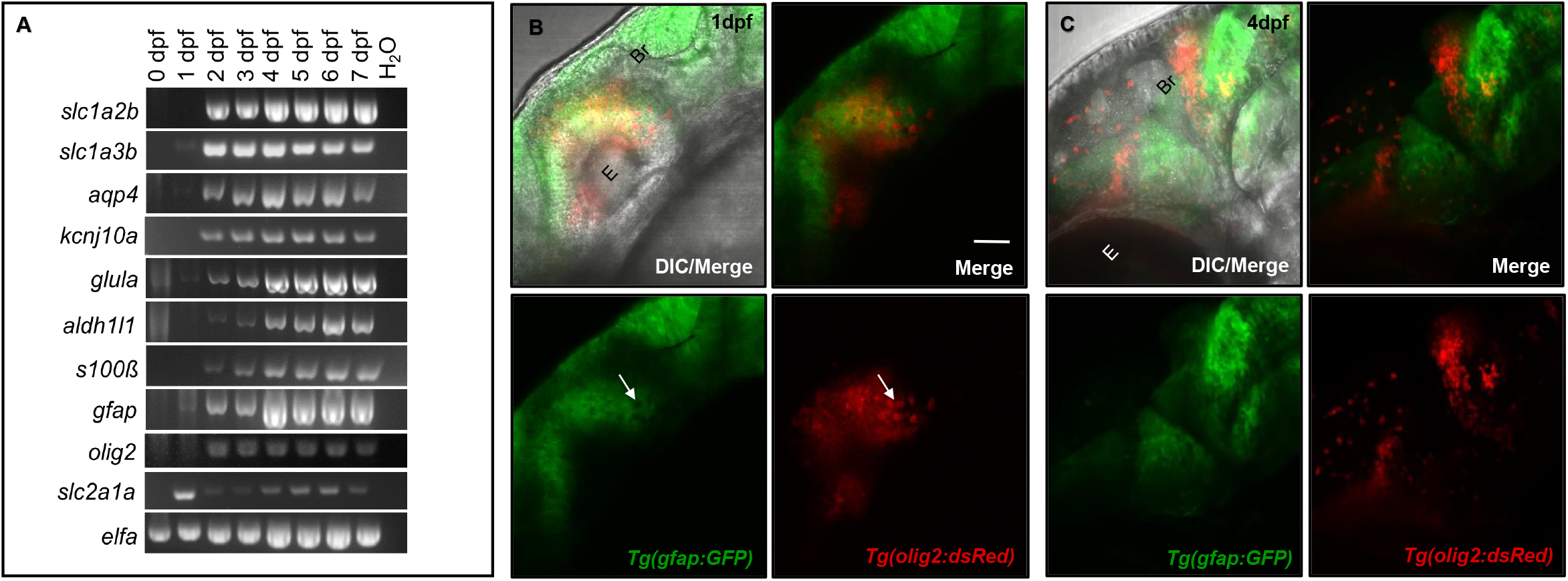
Glial expression increases during zebrafish brain development with distinct lineages. (**A**) RT-PCR panel of different genes commonly expressed in glia, amplified in 0-7dpf zebrafish cDNA samples. A water lane was used as a non-template control. (**B**,**C**) Among the amplicons from panel (**A**), *olig2* and *gfap* label distinct populations at 1dpf (**B**) and 4dpf (**C**) of brain development. E= Eye. Br= Brain. Scale bar= 50µm.

Glia do not uniformly express the same proteins, as they have discrete domains and functions (Freeman, 2010). For example, rodent studies demonstrated the transcription factor Olig2 labeled cells with distinctive lineages for both the myelinating glial progenitors cells (OPCs) and GFAP-negative astrocytes (Tsai et al., 2016; Tatsumi et al., 2018). While spatiotemporally regulated, both sets of glia also interact with the brain vasculature (Sofroniew and Vinters, 2010; Tsai et al., 2016). To first test whether these *olig2* and *gfap* cells have distinct lineages in the zebrafish brain, we crossed *Tg(olig2:dsRed)* animals with the *Tg(gfap:GFP)* line. Live confocal imaging at 1dpf and later at 4dpf revealed these glial cells label distinct populations with no overlap in fluorophore expression (**Figure 1B**,**C**). Therefore, *olig2* and *gfap* cells represent distinct sets of glia in the zebrafish brain for which we could investigate whether they also interact with blood vessels during development.

### Wnt signaling regulates *olig2* migration along vessels and brain endothelial cell expression

As both Olig2 and GFAP glia have relationships with blood vessels in the mammalian brain, we next wanted to determine whether *olig2* and *gfap* glia contacted blood vessels during zebrafish brain development. We first assessed *olig2* glia development with daily live confocal imaging of the *Tg(olig2:dsRed)* line crossed to the vascular line *Tg(fli1a:eGFP)*^*y1*^ from 1dpf to 6dpf (**Figure 2A-F**). Single *olig2* cell populations migrated out of the ventral brain after the onset of CNS angiogenesis (**Supplemental Movie 1**). Furthermore, *olig2* glia co-opted vessels during migration (**Figure2G**) and these interactions increased over 3-6dpf (**Figure 2H**).

**Figure 2.**
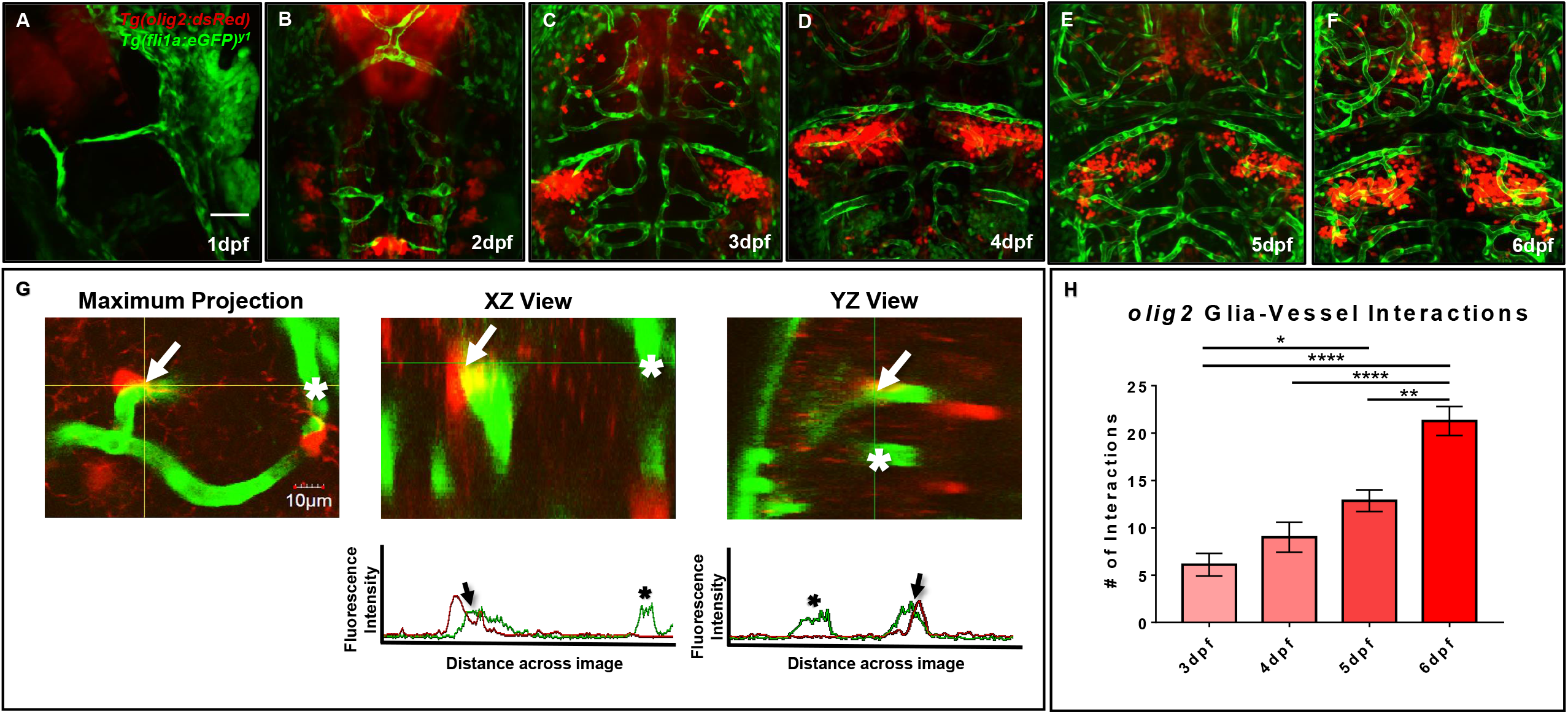
Zebrafish migratory *olig2* glia expand during CNS angiogenesis and co-opt the developing vascular network. (**A-F**) Daily live confocal images of the *Tg(olig2:dsRed)* line (red) crossed to the *Tg(fli1a:eGFP)*^*y1*^ line (green) from 1-6dpf. Red *olig2* glia move out from the ventral brain and migrate along green blood vessels during brain development. Scale bar= 50µm. (**G**) A representative maximum intensity projection of a single *olig2* cell (red) sitting on a vessel (green) (left panel). The orthogonal horizontal (XZ) and vertical (YZ) views (middle and right panel) from the crosshairs where the *olig2* cell sits on a vessel. The orthogonal views are displayed above their corresponding fluorescence line profiles whereby an arrow denotes an *olig2*-vessel interaction and an asterisk shows signal from a nearby vessel without an *olig2* cell. Scale bar= 10µm. (H) *olig2*-vessel interactions quantified from 3-6dpf. A one-way ANOVA with Tukey’s Multiple comparisons test was performed. *p=<0.05, **p=<0.005, ****p=<0.0001. Error bars represent standard error of the mean. n=7-11 per group.

Previous developmental studies in mice demonstrated the necessity of Wnt signaling during Olig2 OPC single cell migration along the brain vasculature (Tsai et al., 2016). To evaluate whether Wnt signaling is a conserved pathway for *olig2* glia-vessel interactions in zebrafish brain, we treated 3dpf *Tg(olig2:dsRed;fli1a:eGFP)* animals by bath application with the Wnt inhibitor XAV939 for 72 hours and performed live confocal imaging at 6dpf. Compared to DMSO vehicle controls, XAV939 treatment significantly reduced the number of *olig2*-vessel interactions after 72 hours (**Figure 3A**). As Wnt signaling is important for both CNS angiogenesis and barriergenesis (Daneman et al., 2009), we teased these processes apart by using the VEGF inhibitor, AV951, during this same time frame of development to solely inhibit new blood vessel formation. Zebrafish Vegf inhibition also reduced *olig2* cell-vessel interactions as would be expected when fewer vessels are present (**Figure 3A**). With these treatments we were also able to dissect whether single *olig2* cell migration was also affected. Quantification of single *olig2* cells during XAV939 and AV951 treatment from 3-6dpf showed that Wnt inhibition and not reduced CNS angiogenesis through VEGF inhibition significantly reduced *olig2* cells (**Figure 3B)** as previously demonstrated in mice (Tsai et al., 2016).

**Figure 3.**
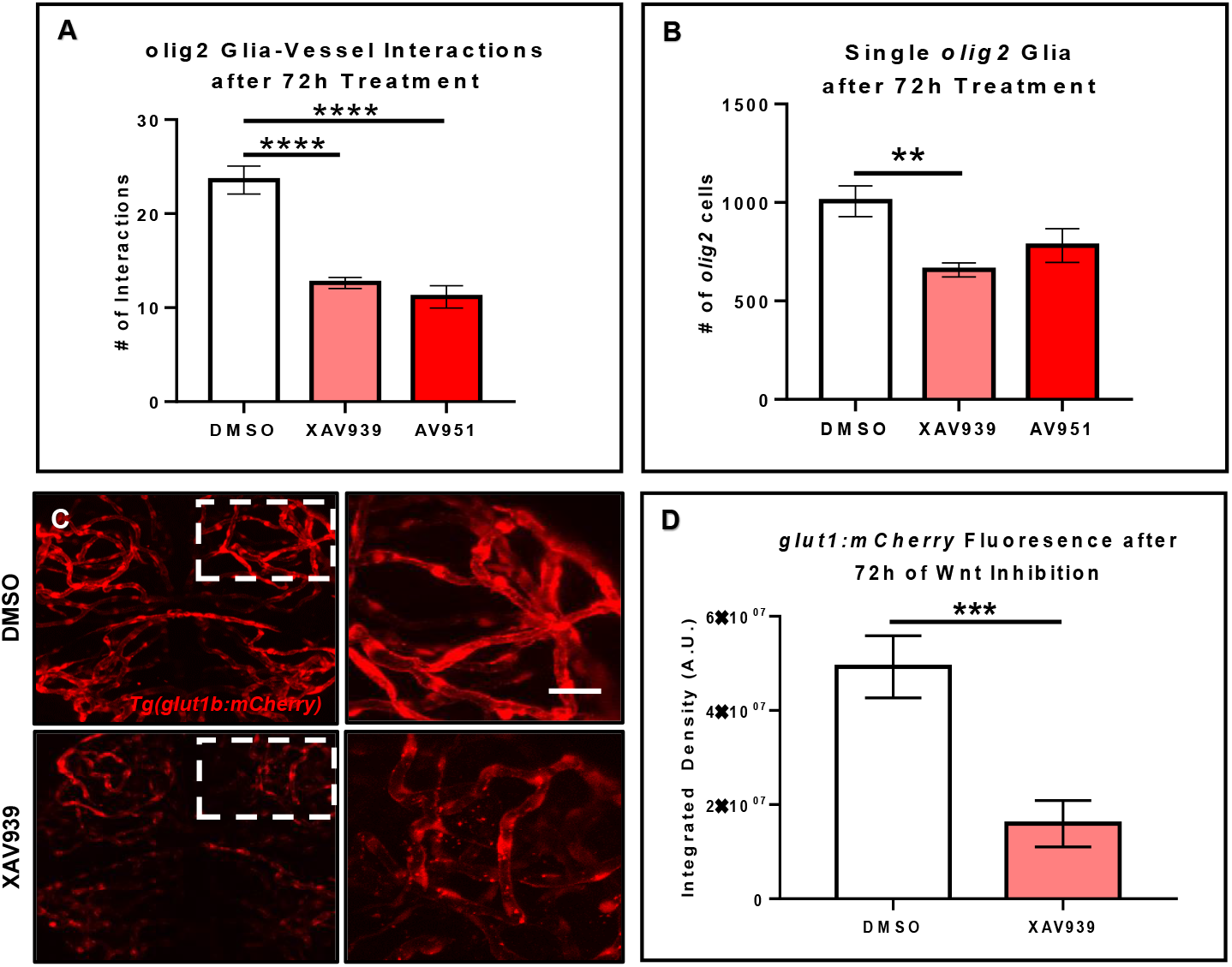
Wnt inhibition decreases *olig2* vessel migration and brain endothelial cell signaling. (**A**) *Tg(olig2:dsRed;fli1a:eGFP)* animals were treated for 72h from 3-6dpf with DMSO vehicle, the Wnt inhibitor, XAV939, or the Vegf inhibitor, AV951. *olig2*-vessel interactions were quantified after 72h of each treatment. A one-way ANOVA with Tukey’s Multiple comparisons test was performed (p<0.0001). Error bars represent standard error of the mean. n= 7-8 animals per group. (**B**) Quantification of individual cells from animals treated in panel (**A**). A one-way ANOVA with Tukey’s Multiple comparisons test was performed (p<0.005). Error bars represent standard error of the mean. n= 10 animals per group. (**C**) Representative live confocal images of *Tg(glut1b:mCherry)* animals treated with either DMSO vehicle control or XAV939 from 3-6dpf. The right panels are the zoomed in areas from the corresponding white boxes in the left panels. Scale bar= 20µm (**D**) Quantification of *glut1b:mCherry* fluorescence after 72h of XAV939 treatment. A two-tailed, unpaired t-test was performed (p<0.001). Error bars represent standard error of the mean. n=9-11 animals per group.

Wnt signaling is known to affect mouse brain blood vessel development as well as protein expression of the BBB protein, GLUT1 (Daneman et al., 2009). Because the reduction in zebrafish *olig2*-vessel interactions was not solely due to a lack of vessels, we asked whether Wnt signaling during this period of CNS angiogenesis also affects brain endothelial cell properties via expression of the BBB marker *glut1b*. To test this, we treated the *Tg(glut1b:mCherry)* line with XAV939 from 3-6dpf, the same time frame of treatment in the *olig2* cell migration experiments. After live confocal imaging and fluorescence analysis, we saw a striking reduction in the *glut1b:mCherry* signal after Wnt inhibition compared to DMSO vehicle control treated siblings (**Figure 3C**). These results demonstrate that *olig2* glia migrate along the zebrafish brain vasculature during development and that this process requires Wnt signaling. Accordingly, Wnt inhibition during 3-6dpf alters brain endothelial cell expression of the BBB transporter *glut1b* and may corroborate why there are fewer *olig2* glia-vessel interactions.

### *gfap* glia have a close association with the developing zebrafish brain vasculature

As we examined how *olig2* glia utilized the brain vasculature as a scaffold during brain development, we next wanted to assess whether zebrafish glia could conversely support vessel growth. In the mammalian brain, GFAP cells outline vessels and cover most of their surface. Consequently, we first asked whether zebrafish *gfap* cells similarly embrace the zebrafish brain vasculature. While we did note a close association of Gfap labeled cells with vessels via immunocytochemistry experiments in the *Tg(flk1:mCherry)* vascular line (**Supplemental Figure 2**), we wanted to more thoroughly follow this line of inquiry with live *in vivo* imaging. We crossed the *Tg(gfap:GFP)* line to the vascular reporter line *Tg(kdrl:mCherry)*, whereby glia are labeled in green and vessels are labeled in red. As a recent study showed zebrafish spinal cord glia mature by 6dpf (Chen et al., 2020), we assessed *gfap*-vessel morphology at 7dpf through multi-photon laser scanning microscopy. Live imaging of 7dpf vessels revealed stunning outlines and points of contact from *gfap* cells as denoted in 3D rendered views and orthogonal projections (**Figure 4**) (n=5 animals, both hemispheres, 20 vessels). These results indicate zebrafish *gfap* glia associate with the developing brain vasculature.

**Figure 4.**
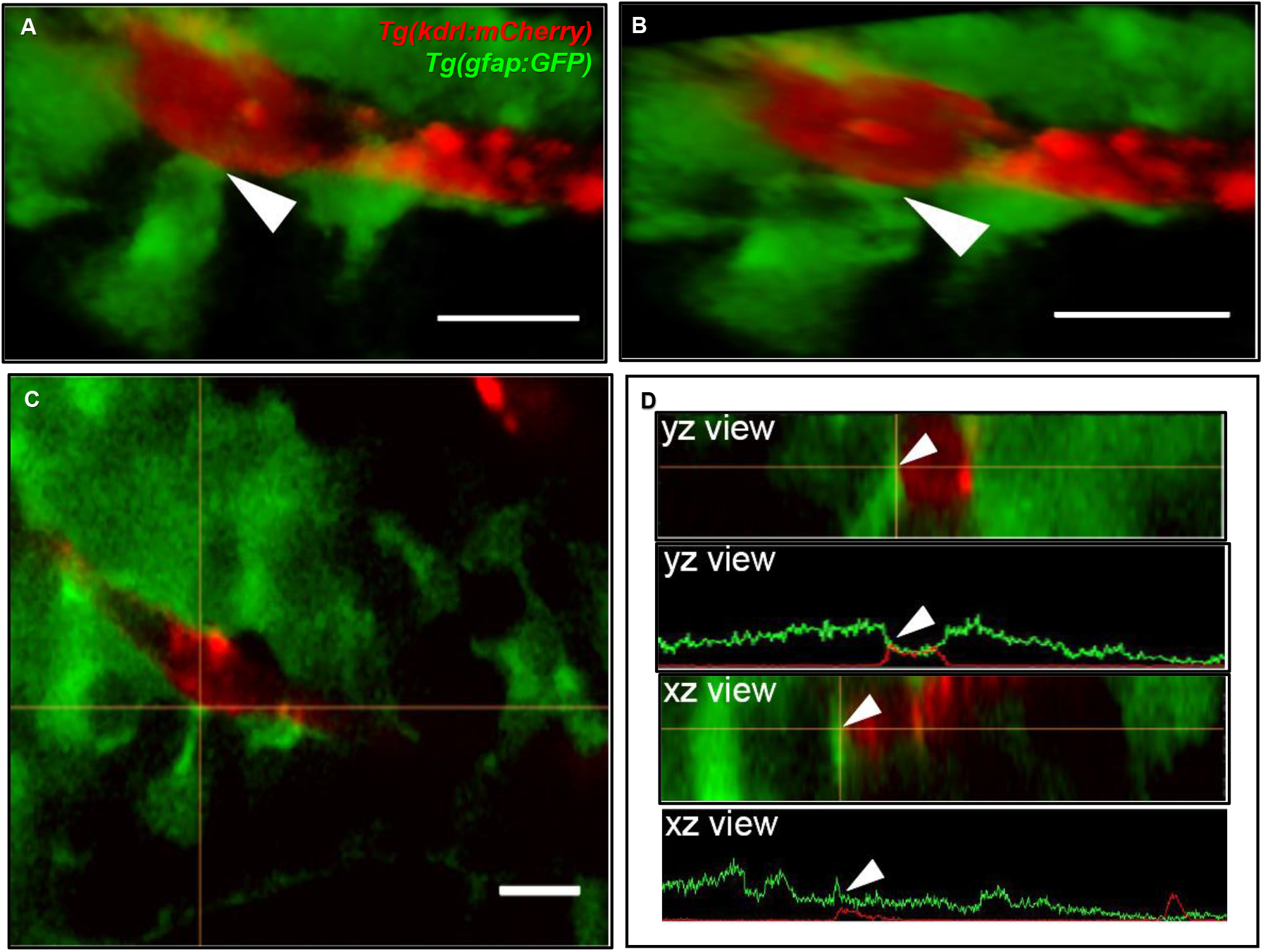
Zebrafish *gfap* glia outline the developing brain vasculature. (**A**) The front-facing view of a volumetric 3D reconstruction showing the interaction (white arrow) of a *gfap* cell process (green) with a brain vessel (red). (**B**) A lateral-view of the same image in panel (**A**). (**C**) A view of the image in panel (**A**) showing the interaction of the green fluorescent signal with the red fluorescent signal at the crosshairs. (**D**) Orthogonal views and line profiles for the vertical (YZ) and horizontal (XZ) crosshairs intersecting (white arrow) where the *gfap* cell meets the vessel. n= 5 animals, both hemispheres, 20 vessels total. Scale bar = 10µm.

### *gfap* glia support brain vessel development via Vegf signaling

We next wanted to examine whether vessel development depended on the presence of *gfap* cells. To determine this, we crossed the *Tg(gfap:nfsB-mCherry)* line to *Tg(fli1:eGFP)*^*y1*^ to assess whether nitroreductase-mediated depletion of *gfap* glia affects vessel structure during CNS angiogenesis. This conditional and inducible cell death occurs when nitroreductase expressed in the cell of interest metabolizes MTZ and generates a cytotoxic metabolite (Curado et al., 2008). We maintained the *Tg(gfap:nfsB-mCherry)* line with heterozygous carriers so we could control for MTZ treated siblings without the *gfap:nfsB-mCherry* transgene (Curado et al., 2008). As Johnson, et al noted brain hemorrhage in 50% of animals after *gfap* ablation from 8-72hpf (Johnson et al., 2016), we used this period to look at vessel morphology and development. Through bright field microscopy, we observed 9% of *gfap:nfsB-mCherry* transgenic animals that were MTZ-treated from 8-72hpf had brain microbleeds (n=33). Live confocal imaging at 3dpf revealed an impairment in vessel growth in MTZ treated *gfap:nfsB-mCherry* transgenic animals (right panel) compared to DMSO and MTZ treated non-transgenic control siblings (left and middle panels) (n= 6-11 animals per group) (**Figure 5A**). While MTZ-treated, *gfap:nfsB-mCherry* transgenic animals had the greatest vessel defects, we did note MTZ-treated non-transgenic siblings had slightly aberrant vessel growth (**Figure 5A**, middle panel). We also performed WMIHC for the apoptotic marker activated Caspase3 in all groups to confirm cell death. 10mM MTZ treatment from 8-72hpf did induce cell death in *gfap:nfsB-mCherry* transgenic larvae as previously reported (Johnson et al., 2016), however we noticed additional 10mM MTZ off-target effects through cell death induced in non-transgenic treated controls compared to DMSO vehicle treated siblings (**Supplemental Figure 3**).

**Figure 5.**
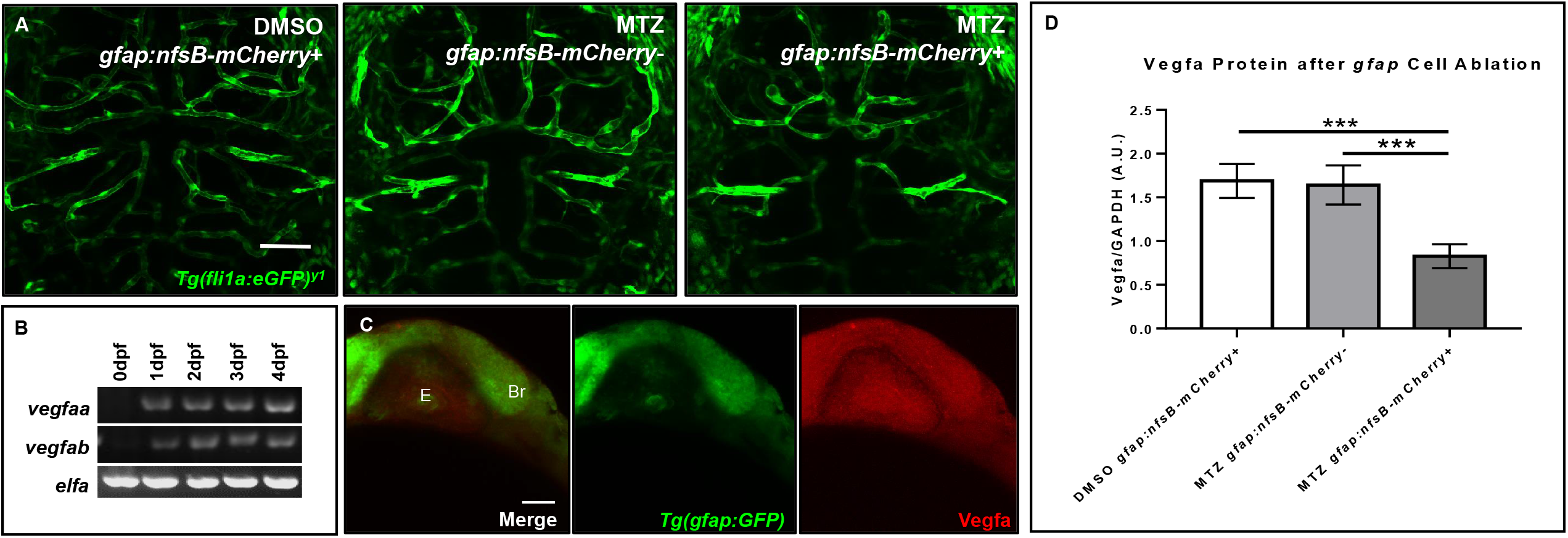
Ablation of *gfap* glia impairs CNS angiogenesis via a reduction in Vegfa. (**A**) Live confocal images of 3dpf animals in the following treatment groups; *gfap:nfsB-mCherry*+ treated with DMSO treated (left), *gfap:nfsB-mCherry*-treated with MTZ (middle), and *gfap:nfsB-mCherry*+ treated with MTZ (right). MTZ treatment in the transgene containing group reduces vessel outgrowth in the brain. n= 6-11 animals per group (**B**) RT-PCR of Vegfa isoforms during 0-4dpf development. (**C**) WMIHC of 1dpf *Tg(gfap:GFP)* (green) stained for Vegfa protein (red). Vegfa is widely expressed during *gfap* cell development. E= Eye. Br= Brain. (**D**) Western blot analysis of Vegfa protein levels in the same treatment groups in (**A**). A two-way ANOVA with Tukey’s Multiple comparisons test was performed (p<0.0005). Error bars represent standard error of the mean. n=5 biological replicates/groups per condition.

We also assessed vessel growth at a later time window when *gfap* cells were ablated from 1-4dpf, starting closer to the onset of CNS angiogenesis. However, we saw less severe vessel impairment through live confocal microscopy at 4dpf, even though 25% of animals (n=25) had some form of microbleed or brain hemorrhage (**Supplemental Figure 4C**). Through live confocal microscopy, we assessed large vascular landmarks such as the posterior communicating segment (PCS) and basal communicating artery (BCA) (**Supplemental Figure 4D-F**) (Isogai et al., 2001). We measured vessel distances between the PCS and connecting BCA vasculature, and found a reduction in the distance between the PCS to the BCA (**Supplemental Figure 4H**). Conversely, inhibition of CNS angiogenesis with VEGF inhibitor AV951 in *Tg(gfap:GFP; kdrl:mCherry)* animals from 1-2dpf did not affect the outgrowth of *gfap* cells, suggesting initial vessel development does not affect *gfap* cell development (**Supplemental Figure 5**).

We next wanted to determine what signaling pathways could be disrupted after *gfap* cell ablation during 8-72hpf when we saw a more dramatic impairment in vessel growth. Taking a candidate approach, we assessed zebrafish Vegf, as it has been shown that astrocytes release VEGF in other highly vascularized tissues like the retina to stabilize growing vessels (Scott et al., 2010). We first evaluated Vegf expression in the developing zebrafish through RT-PCR and WMIHC. Zebrafish express two Vegfa paralogs from 0-4dpf (**Figure 5B**) and the brain and eye ubiquitously express Vegfa protein when *gfap* glia are present at the onset of neural development (**Figure 5C**). Through Western blot analysis, we saw a robust and significant reduction in Vegfa protein after *gfap* cell ablation from 8-72hpf (**Figure 5D**). While we did note off-target Caspase3 activation due to MTZ treatment, this cell death did not cause a robust reduction of Vegfa protein like we saw in MTZ treated *gfap:nfsB-mCherry* transgenic carriers. Therefore, zebrafish *gfap* glia are necessary for proper CNS angiogenesis and this process is partially mediated through Vegf signaling.

### Human glia make contact with the zebrafish brain vasculature

We previously demonstrated in a model of perivascular glioma invasion, that glial-derived tumor cells associate with the developing zebrafish brain vasculature (Umans et al., 2020). However, we wanted to assess whether non-malignant glia would similarly contact zebrafish brain vessels. Cultured human astrocytes were transduced to express GFP and identified by GFAP, GLT-1, Kir4.1, and Aldh1L1 (**Figure6A-E**). We then intracranially implanted these GFP expressing human astrocytes at the midbrain-hindbrain boundary of 6dpf *Tg(kdrl:mCherry)* larvae, whereby blood vessels are labeled with a red fluorophore. 24 hours post implantation, we noted contacts between human glia and zebrafish blood vessels in 60% of the imaged animals (n=8). We followed the same animals 48 hours post-implantation and noted that while fewer human glia were present in each animal compared to images from 24 hours post-implantation (data not shown), 60% of animals displayed glial-vascular contacts between implanted human cells and zebrafish vessels (n=5) (**Figure6F-H**). These results suggest that the attraction to blood vessels is shared between human astrocytes and malignant gliomas, likely mediated by shared signals between endothelial cells, glia, and gliomas. Furthermore, these implantation studies suggest conserved signals support these interactions across vertebrates and thus could be dissected within the genetically tractable zebrafish model.

## Discussion

Our present study adds to a body of glial biology literature by capitalizing on the strengths of the zebrafish model system, whereby we examined glial-vascular interactions in the developing zebrafish brain. Out of the various candidate glial genes we amplified during development, we identified *olig2* and *gfap* cell populations with distinct glial lineages, which is consistent with observations in the mammalian CNS. We also observed that zebrafish *olig2* glia require Wnt signaling to migrate along the vasculature during brain development. In terms of vessel support, *gfap* glia are necessary to stabilize nascent brain vessels through Vegf expression. Furthermore, intracranially implanted human glia reach out to developing zebrafish blood vessels, suggesting conserved signals mediate glial-vascular interactions in zebrafish brain. These results demonstrate the feasibility to dissect endogenous zebrafish glia-blood vessel relationships and the potential of the zebrafish model to uncover novel molecules during vertebrate brain development.

### Wnt signaling during zebrafish *olig2* vessel migration

Of their many roles in the CNS, glia have evolved to make contacts with the cerebral vasculature. These relationships are present among Olig2 OPCs that migrate out of the subventricular zone (SVZ) and onto blood vessels (Tsai et al., 2016) and mature GFAP astrocytes that enwrap the mature BBB (Sofroniew and Vinters, 2010). While we amplified a variety of candidate glial genes during zebrafish CNS angiogenesis (**Figure 1A**), we characterized vascular associations with glia of distinct *olig2* and *gfap* lineages. In the context of the Olig2 cell population, when blood vessels are abolished through loss of Wnt signaling, OPCs fail to migrate out of the SVZ (Tsai et al., 2016). Tsai, H-H. et al demonstrate with a G-protein-coupled receptor (Gpr124) knockout mouse, normal CNS angiogenesis is necessary for proper Olig2 cell migration (Tsai et al., 2016). However, Gpr124 signaling is also important for barriergenesis, and these conclusions may suggest the importance of vessel properties versus the just presence of vessels during OPC migration (Chang et al., 2017; Umans et al., 2017). To assess conservation in the zebrafish and further tease this a part, we established that zebrafish *olig2* cells migrated along the brain vasculature (**Figure 2**). We then added either the Wnt inhibitor XAV939 or the VEGF inhibitor AV951 to *Tg(olig2:dsRed;fli1a:eGFP)* animals from 3-6dpf when we saw a significant increase in *olig2*-vessel interactions (**Figure 2H**) and saw both of these small molecules decreased *olig2*-vessel interactions (**Figure 3A**). Use of these small molecules showed that a significant reduction in single *olig2* cells (**Figure 3B**) is mainly due to Wnt signaling inhibition and not just due to a reduction in vessel growth.

Furthermore, Wnt signaling is important during barriergenesis via the expression of GLUT1 in mammals (Daneman et al., 2009). Because CNS angiogenesis and barriergenesis occur simultaneously in the zebrafish (Umans et al., 2017), we also questioned if Wnt inhibition could regulate zebrafish *glut1b* expression after the onset of barriergenesis from 3-6dpf. Wnt signaling inhibition in *Tg(glut1b:mCherry)* animals showed a dramatic reduction in *glut1b* fluorescence (**Figure 3C**,**D**), indicating a change in brain endothelial cell properties during the same time frame *olig2*-vessel interactions were reduced. These results suggest Wnt inhibition causes *olig2* glia to reduce their migration along vessels that lose BBB proteins, but future studies would need to elucidate if these processes are consequential or mutually exclusive.

### Zebrafish *gfap* glia contact the brain vasculature

Olig2 lineage astrocytes are a distinct subtype from GFAP astrocytes in mouse brain and the zebrafish spinal cord (Park et al., 2007; Tatsumi et al., 2018). Even though GFAP is a hallmark astrocyte label, not all glia express this intermediate filament protein even though it is commonly used to show glial-vascular interactions in mammals (Simard et al., 2003). Therefore, we wanted to evaluate whether zebrafish *gfap* glia have the potential to associate with and stabilize the vasculature in the developing zebrafish brain. Indeed, we demonstrate for the first time in live zebrafish larvae that *gfap* cells have a close relationship with the vasculature in the brain (**Figure 4**). Whether these *gfap* cells are mature astrocytes or radial glia-like cells at this developmental stage remains unknown and still needs to be investigated. Previously, the consensus in the glial biology community was that zebrafish glia were radial-glia like and not bona fide astrocytes (Lyons and Talbot, 2014). However, recent studies indicate morphological maturation of glia in the developing zebrafish brain and spinal cord, suggesting an understudied and exciting biology yet to be explored in the zebrafish model system (Chen et al., 2020). While we did observe 7dpf vessels outlined with *gfap* glia (**Figure 4**), we did not see a complete ensheathment of the vessel wall as it appears in rodent brain (Simard et al., 2003). While we looked at *gfap* glia along vessels during development, Jeong, J et al and colleagues also saw an outline but incomplete covering of Gfap cells around adult zebrafish brain vessels (Jeong et al., 2008). This could be due in part to the use of antibodies, as studies with organotypic slice culture have demonstrated that GFAP antibody labeling does not fully capture the complex cytoarchitecture of a differentiated astrocyte (Benediktsson et al., 2005). Future zebrafish studies should be aimed at following the lifespan of these *gfap* glia into adulthood with the use of transgenic cell membrane labeling.

### *gfap* zebrafish glia stabilize nascent vessels through Vegf

Moreover, through this study we demonstrate the necessity of zebrafish *gfap* cells for brain blood vessel development. Larvae with *gfap* cells ablated from 8-72hpf had impaired CNS angiogenesis and areas with collapsed vessels (**Figure 5A**, right panel). Similar to other mutants that exhibit abnormal vessel patterning, *Tg(gfap:nfsB-mCherry)* MTZ treated animals displayed edema around the pericardium and yolk sac (Roman et al., 2002) (data not shown). As we determined through Western blot analysis, *gfap* ablated animals have a significant reduction in Vegfa protein (**Figure 5D**). The Vegfa isoform is important for vessel dilation and could be why we saw collapsed vessels in our MTZ treated transgenic animals (Nakatsu et al., 2003). A similar study demonstrated zebrafish *gfap* cells modulate vascular patterning in the spinal cord through Vegf signaling (Matsuoka et al., 2016). A reduction in mouse VEGF-A also disrupts vascular plexus patterning in the developing brain and GFAP^-/-^ mice have poorly vascularized white matter (Liedtke et al., 1996; Haigh et al., 2003). Regardless if zebrafish *gfap* glia are more radial glia-like versus a mature astrocyte, immature radial glia help stabilize the vasculature during development and mature astrocytes maintain the BBB (Ma et al., 2013; Heithoff et al., 2020).

A limitation of this *gfap* cell ablation model is that it utilized the nitroreductase system in which we do not get a complete ablation of *gfap* cells. Additionally, we noticed some off-target effects of MTZ on vessel development in our transgene negative treated control siblings (**Figure 5A**, middle panel), yet this biology was robustly worsened in MTZ treated animals with the *gfap:nfsB-mCherry* transgene (**Figure 5A**, right panel). Interestingly, unlike a previous study that dechorionated 8hpf embryos prior to MTZ treatment, we did not perform this and still saw biological effects. An additional limitation of our study was not all MTZ treated carriers of the *gfap:nfsB-mCherry* transgene were homozygous. This could be why we did not see as high of a brain hemorrhage percentage in animals compared to the study performed by Johnson and colleagues whereby they saw 50% hemorrhage after 8-72hpf MTZ treatment (Johnson et al., 2016). While brain hemorrhage was not 100% penetrant in the animals they treated, we noted a similar rate, whereby 25% of animals would be homozygous carriers and close to half of that percentage (9%, 33 animals) had brain hemorrhage after treatment. We chose to maintain heterozygous transgene crosses so we could incorporate an important control for MTZ treatment in non-transgenic siblings (Curado et al., 2008) as MTZ has been shown to have some neurotoxicity in human brain (Kuriyama et al., 2011). These potential off-target effects could also be why we saw substantial Caspase3 staining in non-transgenic MTZ treated control siblings, although it was not statistically significant from DMSO vehicle treated siblings (**Supplemental Figure 3D**).

### Glial cell diversity and signaling across species

Significant glial biology has been uncovered with a breadth of model organisms. The field has traditionally utilized mammals to study glial cells but other model systems have emerged such as *C. elegans* (Oikonomou and Shaham, 2011), *Drosophila melanogaster* (Freeman, 2015), and zebrafish (Johnson et al., 2016; Chen et al., 2020). Zebrafish are vertebrates that possess high genetic conservation to mammals and are suitable for live imaging experiments during development, but their glial identity has been controversial as to whether they possess bona fide astrocytes (Lyons and Talbot, 2014; Muñoz-Ballester et al., 2020). However, a recent study indicated that larval zebrafish possess similar glial proteins, morphology, and biological responses as mature mammalian astrocytes (Chen et al., 2020). Furthermore, a recent toxicity study showed Aflatoxin B1 was toxic to human astrocytes *in vitro* and to the development of glial cells in zebrafish *in vivo* (Park et al., 2019). It is also possible that differences between mammalian and zebrafish glia have redundant roles based on evolutionary divergence, but this is not yet fully understood.

A better understanding of glial cell diversity in multiple model organisms would further elucidate the roles glia play during brain development, CNS homeostasis, BBB maturation, and their importance from an evolutionary perspective (Bundgaard and Abbott, 2008). Zebrafish are advantageous for use in genetic and developmental studies, yet their glial cell maturity is still actively in question (Muñoz-Ballester et al., 2020). While there are differences between mammals and zebrafish, this diversity may enhance our overall understanding of biological processes. For example, zebrafish retain a regenerative capacity that is not present in mammals and the differences in these glial cells may give clues as to why the human CNS does not have this function (Goldshmit et al., 2012). Therefore, more detailed zebrafish studies will answer questions that remain regarding the model system while simultaneously dissecting glial biology.

GFAP is one of the most popular markers to label mature astrocytes, but it is does not uniformly label all healthy cells and is a robust marker of reactive gliosis (Sofroniew and Vinters, 2010). Radial glia also express GFAP during development but mature GFAP human astrocytes later retain specific proteins such as GLT-1 after differentiation (Sofroniew and Vinters, 2010; Roybon et al., 2013). While various astrocyte preparations have their strengths and limitations regarding protein expression (Lange et al., 2012), we wanted to demonstrate cultured human cells were differentiated and whether they would interact with zebrafish brain vessels. Our human astrocyte cultures expressed GFAP, possibly due to being cultured with media containing serum. However, these cultured astrocytes expressed additional proteins that would deem them differentiated, such as Kir4.1, GLT-1, and ALDH1L1 (**Figure 6B-E**). Expression of these markers was important to denote because a mature astrocyte may utilize similar pathways to make contact with the vasculature in the developing and mature brain. We chose to inject 6dpf animals as Chen and colleagues recently demonstrated the maturity of zebrafish spinal cord glia around this time point (Chen et al., 2020). Unlike tumor cells that are mis-programmed to sustain uncontrolled cell growth, we noted that 6dpf animals implanted with cultured astrocytes had fewer cells 48 hours post-implantation compared to 24 hours post-implantation (data not shown). This could be due in part to the environment of a developing brain as well as the compact space in which the human cells had to survive. It is possible that normal astrocytes cannot shrink their cell size as glial derived tumors can (Watkins and Sontheimer, 2011). If our cultured cells were reactive, they may also not survive because that function is not compatible inside a developing brain. As these human astrocytes made contacts with the zebrafish vasculature (**Figure 6F-H**), this suggests that mechanisms that support survival of human astrocytes in the developing zebrafish brain differ from the mechanisms that establish glial-vascular interactions. Nonetheless, because these human cells made close contact with zebrafish vessels, these experiments substantiate studying endogenous glial-vascular interactions in the developing zebrafish brain.

**Figure 6.**
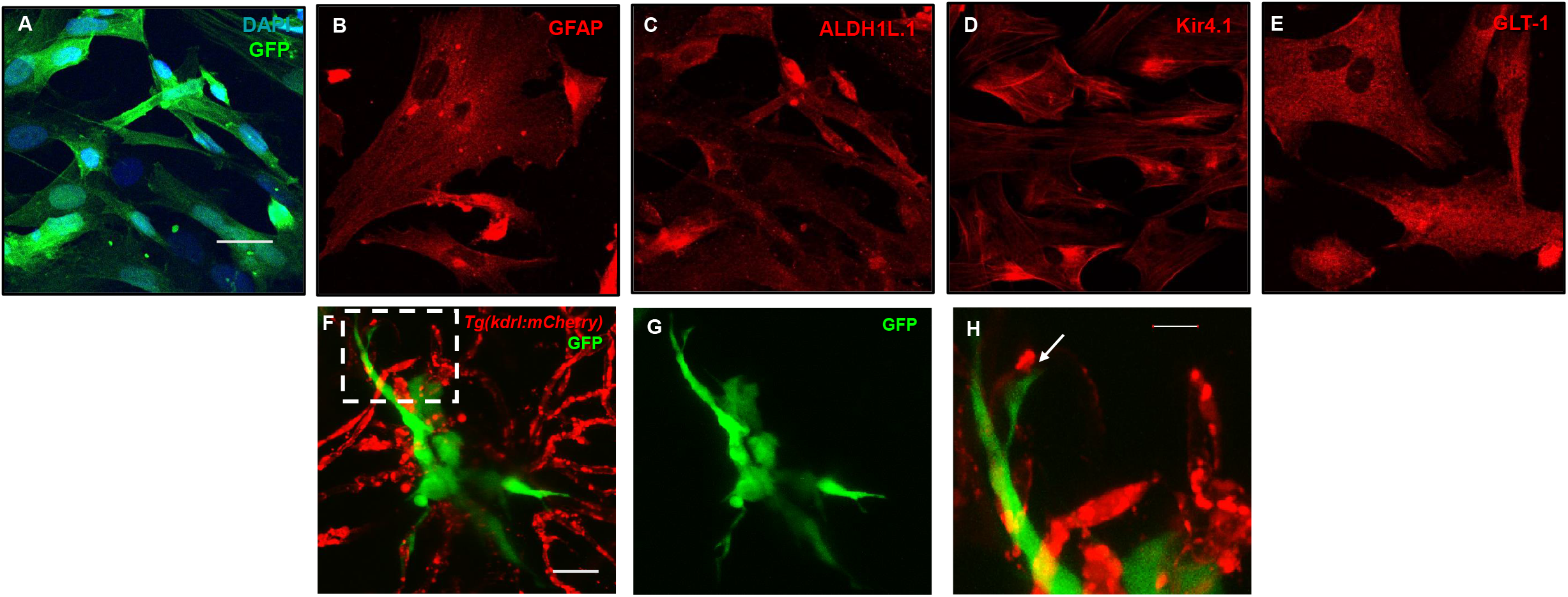
Mature human astrocytes contact developing zebrafish brain vessels. (**A-E**) Immunocytochemistry of GFP-expressing human astrocytes (green), with DAPI counterstained nuclei (blue) (**A**), the common marker GFAP (**B**), and mature markers ALDH1L1 (**C**), Kir4.1 (**D**), and GLT-1 (**E**) (red). Scale bar = 30µm. (**F**) 48 hours post-implantation, human astrocytes (green) reach out to the surrounding brain vasculature (red) of a 6dpf *Tg(kdrl:mCherry)* zebrafish larvae. (**G)** Green channel highlighting the astrocyte’s morphology. (**H**) A zoomed in view from the white dotted box in panel (**F**) showing the astrocyte reaching out to a vessel. n=5 animals. Scale bar = 10µm.

### Future Directions

Developmental biology provides a framework for the mechanisms required to generate healthy tissues and these pathways typically are dysregulated in disease. For example, gliomas possess invasive *olig2* cell populations that migrate along blood vessels during tumor expansion just as *olig2* OPCs migrate during brain development (Tsai et al., 2016; Griveau et al., 2018). Furthermore, glial-vessel relationships hold great importance as cerebral vascular health is tightly linked to neurological diseases. For example, *Gfap*-mediated deletion of *Cerebral cavernous malformation 3* (*Ccm3*) in mice demonstrate cell non-autonomous effects on vascular development similar to Cerebral cavernous malformation disease in humans (Louvi et al., 2011). Additionally, a recent study with a 16p11.2 deletion autism syndrome mouse model displays cerebral angiogenesis impairments that lead to future neurovascular detriments (Ouellette et al., 2020). While Ouellette and colleagues did not see any changes in astrocytes surrounding postnatal and adult blood vessels, it would be interesting to look at embryonic stages and assess the radial glial that may stabilize the nascent vessels that develop abnormally. Therefore our developmental biology studies raise an interesting and open-ended question; are there common pathways in developing glia that stabilize nascent vessels and differentiated glia that maintain mature vessels in the healthy brain? While our studies were limited to a small portion of amplified glial genes and the *olig2* and *gfap* transgenic lines, an array of glia have yet to be studied in zebrafish (**Supplemental Table 1**) and novel genes are likely to be discovered. Taking advantage of pharmacological studies, live imaging, fast development, and genetic manipulation studies over short periods of time, we demonstrate the feasibility and attractiveness of using zebrafish to study glial-vessel interactions. Therefore, additional zebrafish studies could clarify glial diversity in the CNS, elucidate novel mechanisms important for CNS and vascular development, and illuminate the associated molecular cues vital for glial-vascular biology.

## Author Contributions

R.A.U conceived the study and experimental design, acquired and analyzed data, and drafted the manuscript. C.P. aided in experimental design, acquired, and analyzed data. W. A.M. III aided in experimental design, acquired, and analyzed data, and H.S. aided in experimental design and edited the manuscript.

## Acknowledgments

We would like to thank Drs. Michael Taylor, Sarah Kucenas, Wilson Clements, and Chris Lassiter for providing us with zebrafish lines that made this study possible. Thank you to Mary Grace Milam for her assistance getting these FBRI zebrafish projects started. We would also like to thank members of the Pan laboratory for experimental feedback, the Caspase-3 antibody/protocol, and use of their stereomicroscope.

