## Supplemental Figures and Legends for "Using zebrafish to elucidate glial-vascular interactions during CNS development"

#### Supplemental Figure 1

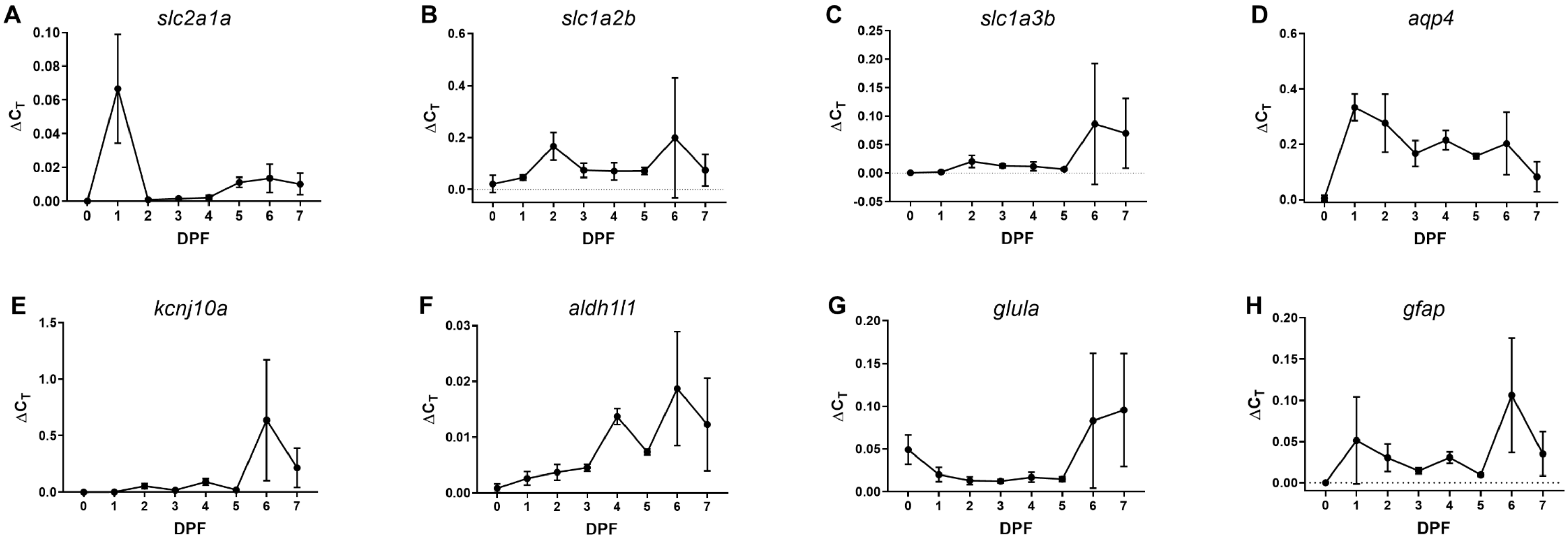

### Supplemental Figure 2

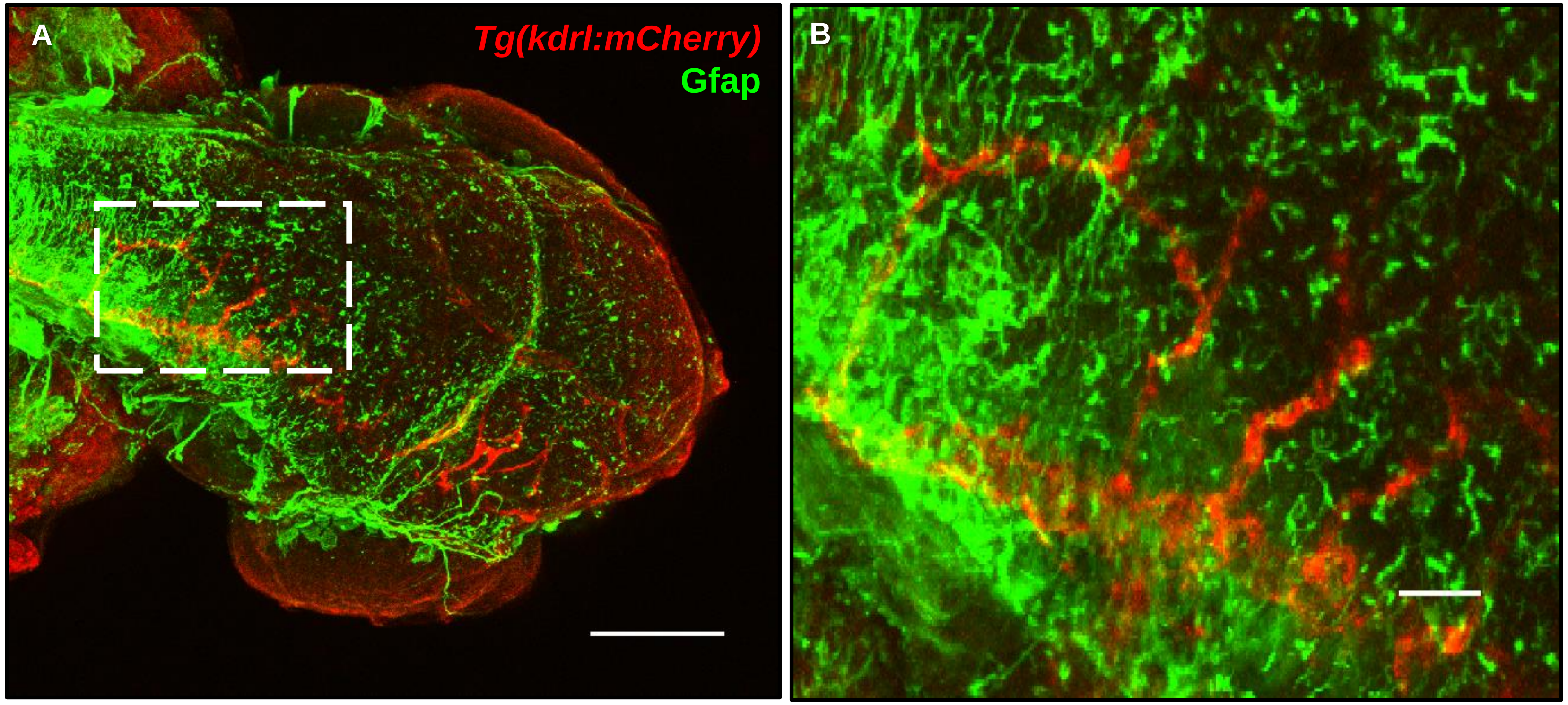

### Supplemental Figure 3

C

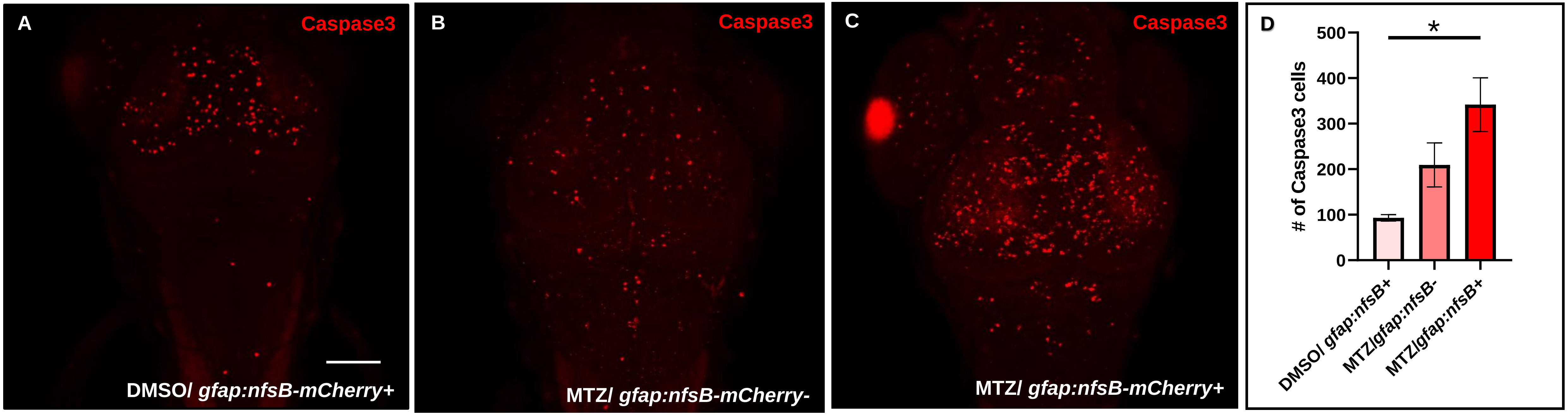

**Supplemental Figure 4**

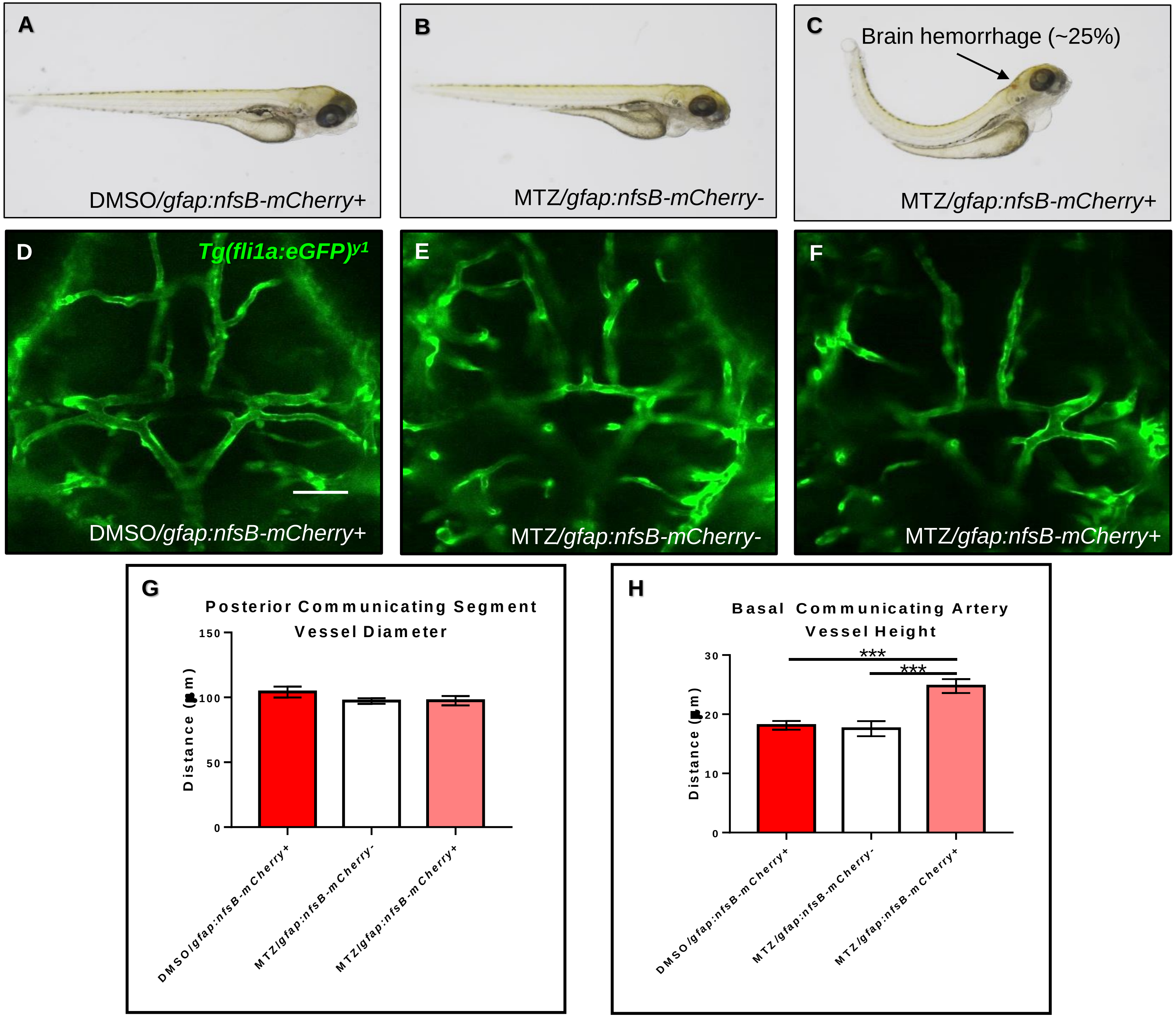

#### Supplemental Figure 5

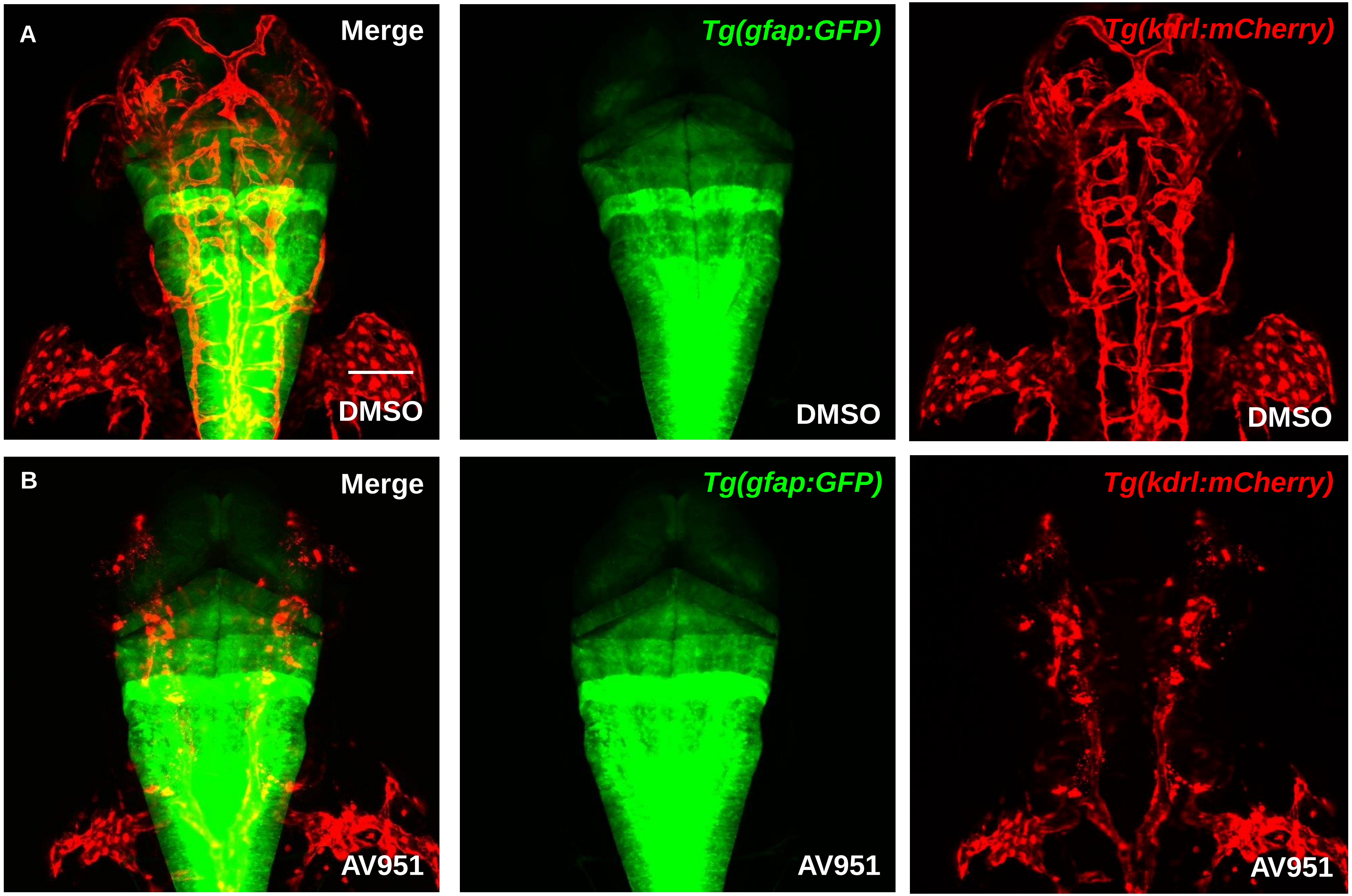
